# Growth cone advance requires EB1 as revealed by genomic replacement with a light-sensitive variant

**DOI:** 10.1101/2022.09.22.509085

**Authors:** Alessandro Dema, Rabab Charafeddine, Shima Rahgozar, Jeffrey van Haren, Torsten Wittmann

## Abstract

A challenge in analyzing dynamic intracellular cell biological processes is the dearth of methodologies that are sufficiently fast and specific to perturb intracellular protein activities. We previously developed a light-sensitive variant of the microtubule plus end tracking protein EB1 by inserting a blue light-controlled protein dimerization module between functional domains. Here we describe an advanced method to replace endogenous EB1 with this light-sensitive variant in a single genome editing step enabling this approach in human induced pluripotent stem cells and derived neurons. We demonstrate that acute and local optogenetic EB1 inactivation in developing cortical neurons induces microtubule depolymerization in the growth cone periphery and subsequent neurite retraction. In addition, advancing growth cones are repelled from areas of blue light exposure. These phenotypes were independent of the neuronal EB1 homolog EB3, revealing a direct dynamic role of EB1-mediated microtubule plus end interactions in neuron morphogenesis and neurite guidance.

## Introduction

Microtubules (MTs) are essential components of the cytoskeleton in both mature neurons, in which polarized MT tracks support axonal transport, as well as during nervous system development (Kapitein and Hoogenraad, 2015; van de Willige et al., 2016; Atkins et al., 2022). During neuron differentiation, expression of many microtubule-associated proteins (MAPs) is upregulated, and numerous neurodevelopmental diseases are being linked to genetic alterations in MAPs, MT motors and tubulin itself (Franker and Hoogenraad, 2013; Maillard et al., 2022). Developing neurites must elongate over long distances integrating extrinsic cues to achieve correct nervous system topology. Neurite growth and pathfinding is driven by adhesion and F-actin dynamics in the growth cone, the advancing terminal structure of growing neurites. Analogous to the role of MTs in migrating cells, dynamic MTs enter the F-actin rich growth cone periphery and these so-called ‘pioneer’ MTs participate in growth cone advance and guidance downstream of extracellular cues (Vitriol and Zheng, 2012; Liu and Dwyer, 2014).

+TIP proteins that associate with growing MT plus ends, such as spectraplakins (MACF1), the adenomatous polyposis coli protein APC, CLASPs and neuron navigator proteins (Nav1) play critical roles in neuron morphogenesis (Coles and Bradke, 2015; Cammarata et al., 2016). For example, Nav1 mediates interactions between dynamic MT plus ends and F-actin filaments (Sánchez-Huertas et al., 2020) and CLASPs are involved in cell-matrix adhesion site remodeling (Stehbens et al., 2014). However, the role of +TIPs in neuron morphogenesis is inferred mostly from genetic knockout phenotypes in several models systems from different species. It has not been tested how +TIP association with growing MT ends participates in controlling growth cone dynamics in real time in developing human neurons in part due to an absence of technology to acutely inhibit these interactions. All known +TIPs bind to growing MT plus ends through small adaptor proteins of the end-binding (EB) family, encoded in humans by three MAPRE genes. Here, we utilize our recent optogenetic tool to disrupt EB1 function and thus +TIP MT-binding in growth cones of human induced pluripotent stem cell (hiPSC)-derived developing cortical neurons. In addition to demonstrating a novel strategy to generate photosensitive variants of multidomain proteins in a single genome editing step, we show that EB1 is necessary to stabilize MTs in the growth cone periphery and to maintain growth cone advance.

## Results and discussion

### One-step genome editing to generate photosensitive protein variants

We recently reported an optogenetic system to inactivate end-binding protein 1 (EB1) with high spatial and temporal accuracy in living cells by inserting a light-sensitive LOV2/Zdk1 dimerization module between the N-terminal +TIP and C-terminal MT-binding domains of EB1 (Fig. 1A) and demonstrated effects on MT dynamics and function in interphase and mitotic human cancer cells (Dema et al., 2022a; van Haren et al., 2018). However, our initial method of generating π-EB1 cell lines required sequential genetic knockouts and re-expression of photosensitive EB1 variants. To enable and demonstrate the utility of this LOV2/Zdk1-mediated multidomain splitting strategy to photoinactivate proteins expressed at endogenous levels in more complex cell systems such as an hiPSC model of neuronal morphogenesis, we devised a CRISPR/Cas9-mediated genome editing strategy to directly insert the LOV2/Zdk1 dimerization module into the EB1 gene and thus convert endogenous EB1 into the photosensitive π-EB1 variant in one genome editing step.

**Figure 1.**
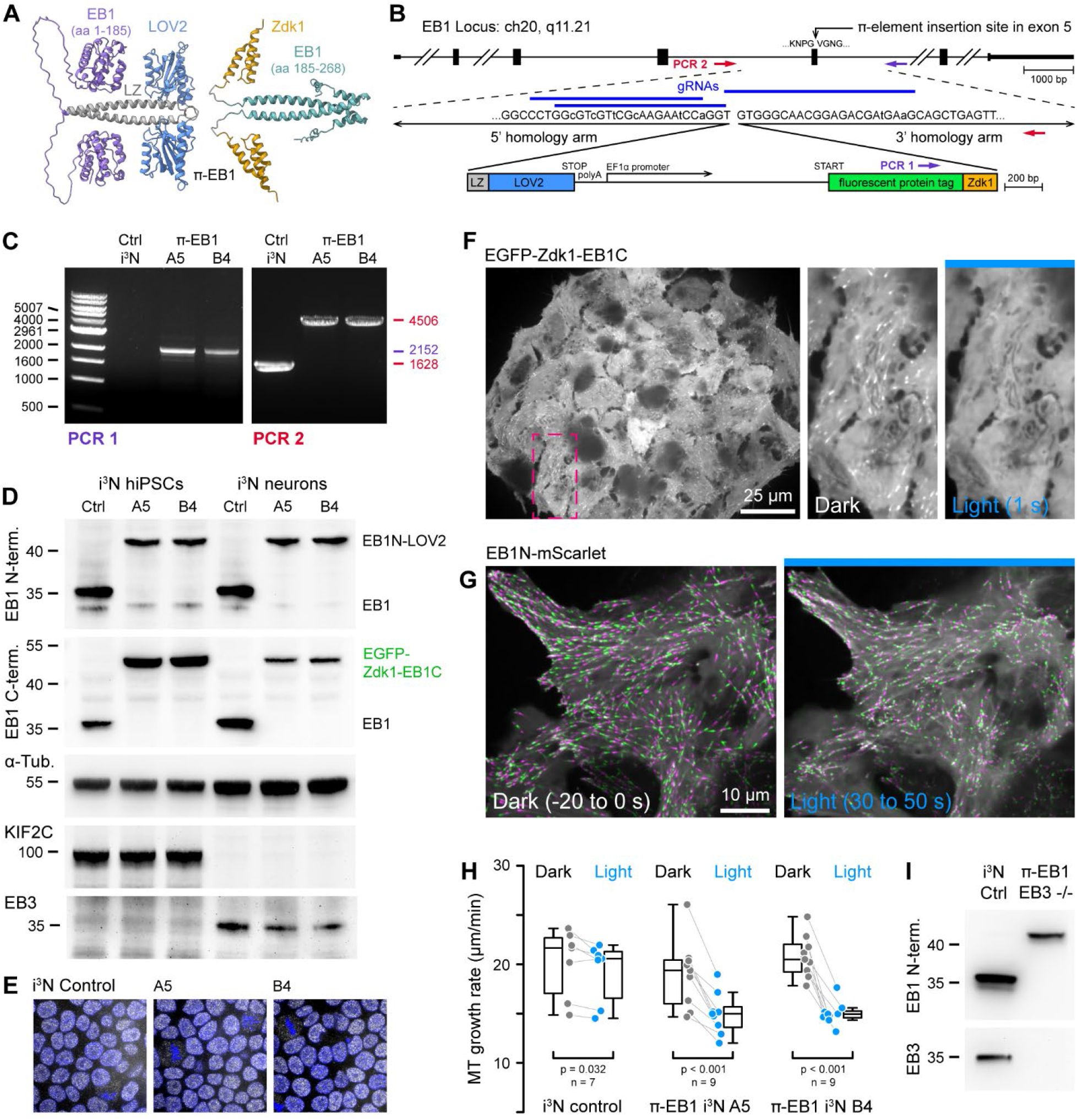
One-step genome editing to replace EB1 with a photo-sensitive variant. **(A)** AlphaFold2 model of the π-EB1 tetramer (Mirdita et al., 2022). Note that AlphaFold2 does not correctly predict relative domain positions and did not capture the LOV2/Zdk1 interaction correctly although a structure of the LOV2/Zdk1 dimer has previously been determined (Wang et al., 2016). **(B)** Overview of the one-step CRISPR/Cas9-mediated insertion of a π-element construct containing the photosensitive LOV2/Zdk1 module, a fluorescent protein marker and an internal EF1α promoter. Arrows indicate location of PCR primers. Lowercase letters indicate mutations introduced to make the HDR template resistant to Cas9 cleavage. **(C)** Genomic PCR to validate π-element integration into the endogenous EB1 locus. The two clones shown are homozygous as there is no short product in PCR2. **(D)** Immunoblots of control (Ctrl) and π-EB1 i^3^N clones before and after 2 days of neuron differentiation with antibodies as indicated showing replacement of EB1 by the photosensitive π-EB1 variant and expected +TIP expression level changes associated with neuron differentiation. **(E)** Comparison of nuclear Oct4 staining (white) as a pluripotency marker in control and π-EB1 i^3^N hiPSC colonies. Nuclei are identified with DAPI (blue). **(F)** Image of a π-EB1 i^3^N hiPSCs colony with magnified images on the right showing dissociation of EGFP-Zdk1-EB1C from growing MT ends in blue light. **(G)** π-EB1 i^3^N hiPSCs transiently expressing an mScarlet-tagged EB1N MT-binding domain before and during blue light exposure. Maximum intensity projections in alternating colors over 20 s at 3-s intervals illustrate attenuation of MT growth during blue light exposure. **(H)** Quantification of the median MT growth rate per cell before and during blue light exposure in control and π-EB1 i^3^N hiPSCs. **(I)** Immunoblot of control and π-EB1 and EB3-/-i^3^N neurons showing expression of π-EB1 and deletion of both EB1 and EB3.

Evaluating different designs of what we named π-elements in H1299 human cancer cells, we found that homozygous integration of a π-element in which an internal EF1α promoter drives expression of the C-terminal π-EB1 half while the N-terminal half remains under control of the endogenous promoter resulted in the most balanced expression of both π-EB1 parts and yielded EB1 protein levels comparable to control cells (Fig. S1; Fig. 1B). In contrast, an internal ribosome entry site (IRES) resulted in very poor expression of the C-terminal half. Of note, self-cleaving 2A peptides also did not work because the C-terminus of the LOV2 domain cannot be modified without greatly inhibiting Zdk1 binding (Wang et al., 2016).

In i^3^N cells, an hiPSC line that expresses Ngn2 under a doxycycline-induced promoter to allow inducible differentiation into cortical glutamatergic neurons (Wang et al., 2017), homozygous π-element integration into both copies of exon 5 of the endogenous MAPRE1 (EB1) gene was validated by genomic PCR and immunoblot (Fig. 1 C,D), and did neither alter the stem cell properties of i^3^N cells (Fig. 1E) nor affect the Ngn2-driven neuronal differentiation program as indicated by expected changes in expression levels of marker proteins (Fig. 1D) such as EB3 and KIF2C that increase or decrease, respectively, during neuronal differentiation (Fig. S2A-C) (Blair et al., 2017). To visualize π-EB1 photodissociation in genome edited i^3^N cells, we chose to tag the π-EB1 C-terminal part with EGFP so that longer wavelengths imaging channels remain available. Clonal π-EB1 i^3^N hiPSC colonies homogenously expressed EGFP-Zdk1-EB1C, and as expected, EGFP-Zdk1-EB1C rapidly dissociated from growing MT ends during blue light exposure in both i^3^N stem cells (Fig. 1F) and differentiating neurons (Fig. S2D). In addition and similar to what we previously observed in human cancer cells (van Haren et al., 2018), π-EB1 photoinactivation significantly attenuated MT growth in i^3^N hiPSCs transiently expressing an mScarlet-tagged EB1 N-terminal construct that remains on MT ends to track MT growth during blue light exposure (Fig. 1F,G; Video 1). Because EB3 is predominantly expressed in neurons and EB1 and EB3 are thought to be at least partially functionally redundant, we also knocked out the EB3 gene in π-EB1 i^3^N hiPSCs (Fig. 1I; Fig. S3A-C) to be able to ask how EB1 contributes to neuron morphogenesis and compare the relative contributions of EB1 and EB3.

### EB1 stabilizes growth cone microtubules

Upon Ngn2-induced differentiation, π-EB1 EB3-/-i^3^N cells sprouted neurites normally (Fig. S3D; Video 2), and as expected, the EGFP-tagged C-terminal half of π-EB1 associated with growing MT ends in neurites with the majority of growing MT ends localized to growth cones. Of note, because MT growth is slower in neurons than in proliferating cells, EGFP-Zdk1-EB1C comets were less elongated in developing neurons and appeared more punctate (Stepanova et al., 2003). Nevertheless, EGFP-Zdk1-EB1C quickly and reversibly dissociated from MT ends upon blue light exposure in differentiating π-EB1 EB3-/-i^3^N neurons (Figs. 2A, S2D; Video 3).

**Figure 2.**
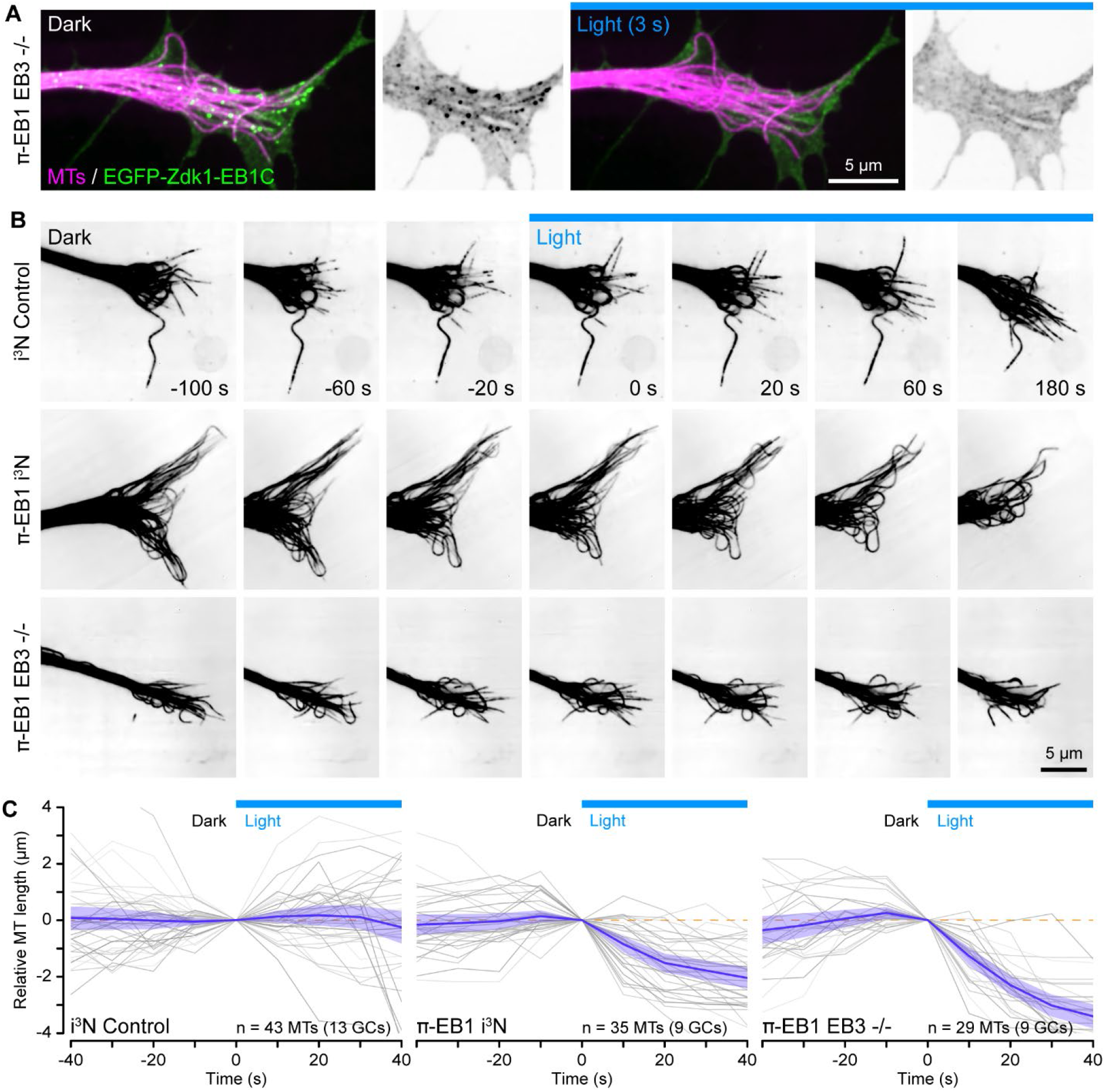
π-EB1 photoinactivation destabilizes growth cone MTs. **(A)** Growth cone of a π-EB1 EB3-/-i^3^N neuron in which MTs were labelled with the far-red cabazitaxel derivative 4-610CP-CTX. To better show individual growth cone MTs, the gamma of the magenta channel was adjusted to 0.6. EGFP-Zdk1-EB1C dissociates from growing MT ends within seconds of blue light exposure. **(B)** MTs labelled with SPY555-tubulin in growth cones of i^3^N neurons with the π-EB1 genotype indicated on the right. Note that MTs continue to dynamically extend into the growth cone periphery in control i^3^N neurons but retract upon blue light exposure in both π-EB1 and π-EB1 EB3-/-i^3^N growth cones. Single channels are shown in inverted contrast for better visibility. **(C)** Quantification of the length change of 3-4 MTs per growth cone the ends of which were clearly visible before and during blue light exposure. Grey lines indicate individual MTs. Blue line is the average of all MTs and the shaded area indicates the 95% confidence interval. The orange dashed line indicates no change.

Using a fluorescently labeled EB1 MT-binding domain, we previously reported in interphase cells and here in proliferating i^3^N hiPSCs that π-EB1 photoinactivation attenuated MT growth. Because we were unable to reliably transfect differentiating i^3^N-derived neurons, we instead used new fluorogenic taxane derivatives as live MT labels (Buceviĉius et al., 2020; Lukinaviĉius et al., 2014) at very low concentrations to evaluate how π-EB1 photoinactivation affected the growth cone MT cytoskeleton. Because taxane binding to intracellular MTs appears slow compared to the MT growth rate (Ettinger et al., 2016), the ends of actively growing MTs are labeled only dimly (Fig. 2A). Thus, these probes are not well-suited to measure MT growth rates but work well to detect MT shortening. In control i^3^N neuron growth cones, MTs dynamically extended from the central domain into the growth cone periphery and the average length of these MTs was not affected by blue light. In contrast, in π-EB1 i^3^N-derived neurons MTs rapidly retracted from the peripheral domain in response to blue light exposure (Fig. 2B,C; Video 4). While this MT retraction was most pronounced in π-EB1 EB3-/-i^3^N neurons, MTs also shortened in π-EB1 i^3^N neurons that still expressed EB3 and in both cases this π-EB1 photoinactivation-induced MT shortening was significantly different from control i^3^N neurons (p < 0.001 by ANOVA and Tukey-Kramer HSD). Because the N-terminal half of π-EB1 that remains on growing MT ends is expressed from the endogenous EB1 promoter at near endogenous levels in the π-EB1 i^3^N neurons and is not overexpressed (Fig. 1D,I), it is unlikely that EB1N-LOV2 interferes with EB3 binding to growing MT ends more than wild-type EB1 although we previously observed that π-EB1 photoinactivation can partially displace EB3 from growing MT ends (van Haren et al., 2018). Thus, these data indicate that EB1 is required for growth cone MT stability and that EB3 cannot rescue EB1-mediated MT growth, consistent with the idea that the function of EB3 differs from that of EB1 and may be more important during later stages of neuron development when EB3 is more highly expressed (Jaworski et al., 2009; Leterrier et al., 2011).

### Acute EB1 inactivation does not immediately change F-actin dynamics

Because ‘pioneer’ MTs that enter the growth cone periphery participate in growth cone guidance (Buck and Zheng, 2002), and MT and f-actin dynamics are coupled both mechanically and biochemically through Rho GTPase signaling, we next asked how π-EB1 photoinactivation affected f-actin dynamics. MT growth is thought to activate Rac1 that drives leading edge actin polymerization, while MT shortening activates RhoA that increases actomyosin contractility through the release of regulatory MT-bound factors (Garcin and Straube, 2019; Wittmann and Waterman-Storer, 2001). To investigate growth cone f-actin dynamics in response to π-EB1 photodissociation-mediated growth cone MT shortening, we used SPY650-FastAct, a fluorogenic jasplakinolide derivative that binds strongly to f-actin filaments and at very low concentrations forms fluorescent speckles that serve as reporters of f-actin flow dynamics (Fig. 3A) (Danuser and Waterman-Storer, 2006). On kymographs perpendicular to the growth cone edge (Fig. 3B), we measured an f-actin retrograde flow rate of 2.8 +/-0.3 µm/min (mean +/-standard deviation) in π-EB1 i^3^N neurons in the dark, similar to previous measurements in growth cones from other vertebrate species (Geraldo and Gordon-Weeks, 2009; Flynn et al., 2012; Gomez and Letourneau, 2014). Although overall growth cone morphology was highly dynamic (Video 5), the growth cone F-actin retrograde flow rate remained remarkably constant during up to 5 minutes of blue light exposure (Fig. 3C). Because retrograde F-actin flow is directly related to the rate of leading-edge actin polymerization, this indicates that EB1-mediated MT growth or +TIP interactions do not immediately influence growth cone actin polymerization dynamics.

**Figure 3.**
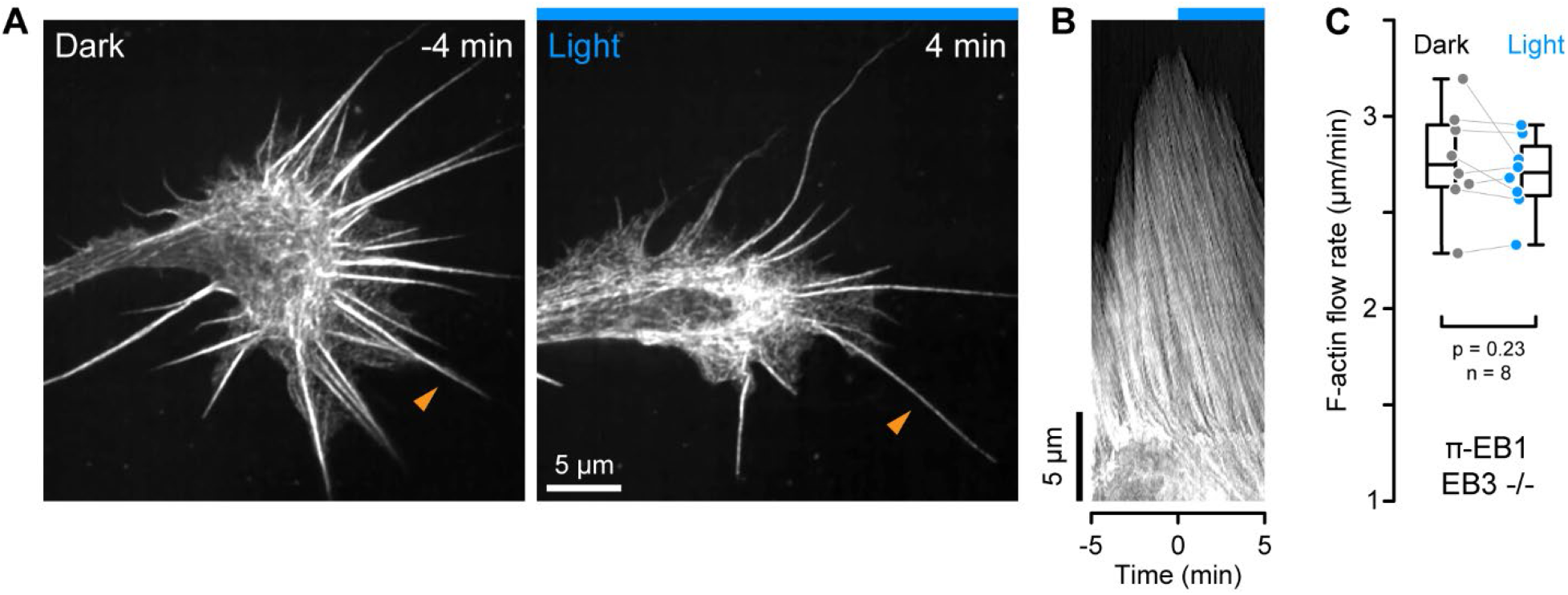
F-actin dynamics in π-EB1 neuron growth cones. **(A)** Growth cone of a π-EB1 EB3-/-i^3^N neuron labelled with SPY650-FastAct before and during blue light exposure. **(B)** Kymograph along the filopodium indicated by orange arrowheads in A illustrating f-actin retrograde flow. Gamma was adjusted to 0.6 to better visualize dim features. **(C)** Quantification of the f-actin retrograde flow rate before and during blue light exposure. Each data point represents the average of 4-6 flow measurements per growth cone.

### EB1 is required for growth cone advance

However, after several minutes of π-EB1 photoinactivation, we noticed that π-EB1 neurites frequently started to retract. Tracking growth cone position over time, >90% of π-EB1 i^3^N neurites retracted in response to blue light over a 15 min observation window (Fig. 4A,B; Video 6), regardless of whether they expressed EB3 or not, again indicating that EB3 is not able to compensate for the acute loss of EB1 function in early neurite development. Similarly, growing π-EB1 i^3^N neurites were unable to cross a blue light barrier in most cases (27 out of 33 cells) sometimes after repeated growth attempts over several hours (Fig. 4C; Video 7). In contrast and as expected, control i^3^N neurons were insensitive to blue light exposure with only 40% of neurites shortening (Fig. 4A,B).

**Figure 4.**
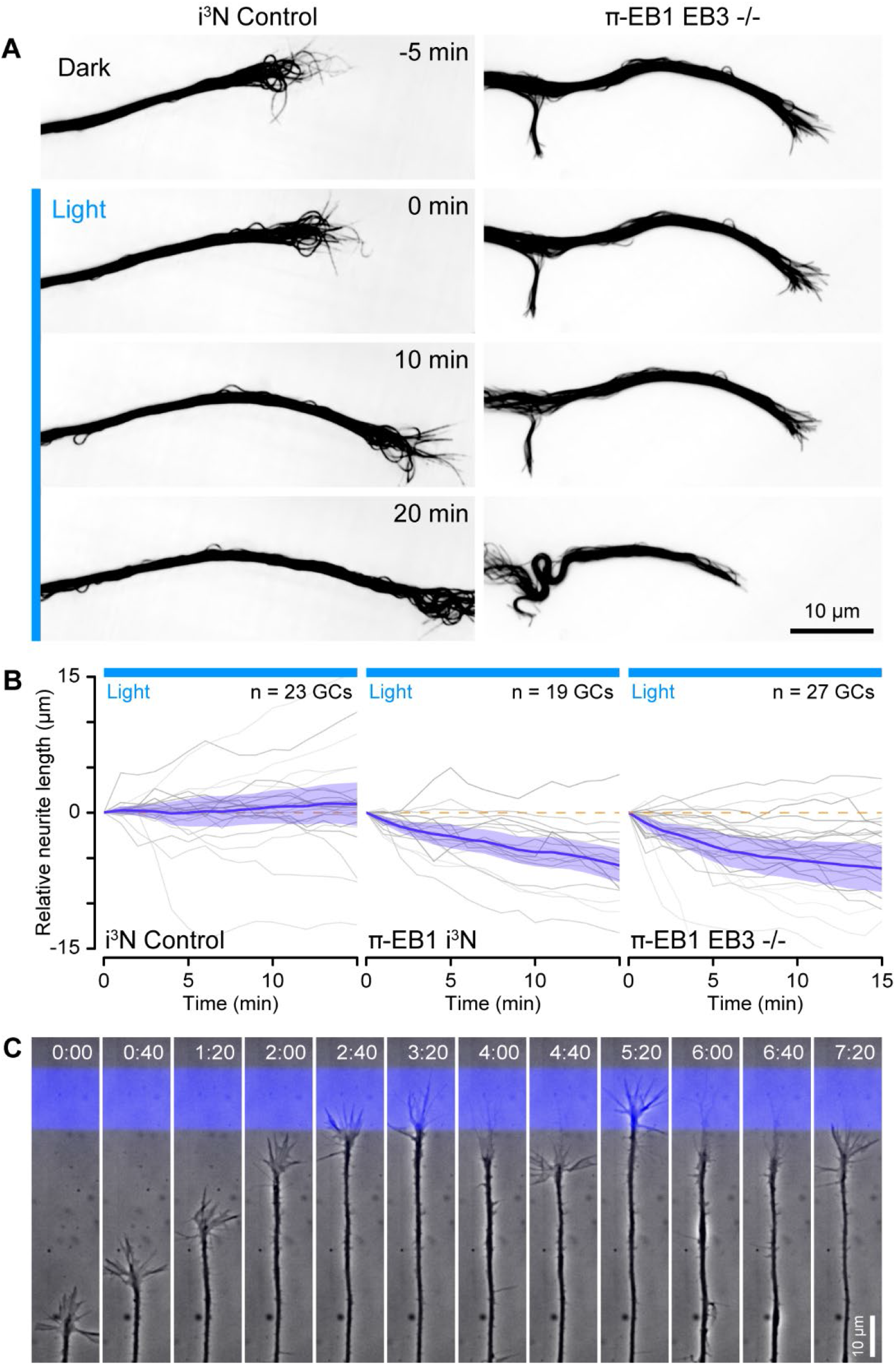
π-EB1 photoinactivation blocks growth cone advance. **(A)** Control and π-EB1 EB3-/-i^3^N neurites in which MTs were labeled with 4-610CP-CTX before and during blue light exposure illustrating retraction of the π-EB1 neurite in blue light while the control neurite continues to advance. **(B)** Quantification of the neurite length change before and during blue light exposure. Grey lines indicate individual neurites. Blue line is the average of all neurites, and the shaded area indicates the 95% confidence interval. The orange dashed line indicates no change. **(C)** Long-term phase contrast time-lapse sequence of a π-EB1 neurite advancing upward on a 10 µm wide stripe of laminin illustrating growth cone retraction every time the growth cone attempts to cross the virtual blue light barrier. Elapsed time is indicated in hours:minutes.

Custom-engineered neuron networks with defined connectivity between individual neurons are a potential key to better understand nervous system information processing (Aebersold et al., 2016), and optogenetic guidance of developing neurites would be a useful tool to build such neuron networks from the bottom up. We therefore tested if more precise blue light exposure to small regions inside growth cones targeting only few MTs could be used to control the direction of growth cone advance (Fig. 5A). However, this experiment was technically very challenging. It was especially difficult to correctly place the blue light exposure ROI such that π-EB1 photoinactivation remained sufficiently spatially restricted. Hence, as a result most neurites (42 out of 58 cells) retracted even with localized blue light exposure. Yet, analyzing GC displacement of neurites that did not immediately retract revealed a turning response away from the blue light exposed area (Fig. 5B).

**Figure 5.**
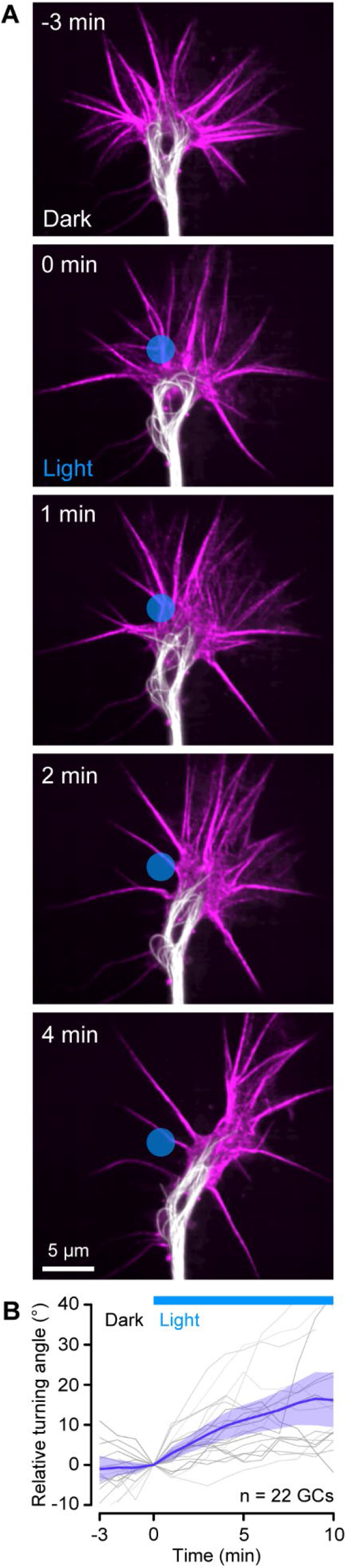
Growth cone turning in response to local π-EB1 photoinactivation. **(A)** Time-lapse of a π-EB1 EB3-/-i^3^N neurite growth cone labeled with SPY555-tubulin (white) and SPY650-FastAct (magenta). The gamma of the tubulin channel was adjusted to 0.6 to better visualize individual growth cone MTs. The blue circle indicates the light-exposed area. **(B)** Quantification of the relative turning angle in response to local blue light exposure. Grey lines indicate individual growth cones. Blue line is the average of all growth cones measurements, and the shaded area indicates the 95% confidence interval.

In summary, we present a novel approach to replace the central MT regulator EB1 with a photosensitive variant by inserting a light-sensitive protein interaction module encoded by a short genetic cassette – that we named π-element – into an inter-domain linker by CRISPR/Cas9-mediated genome editing. It is important to note that in principle the π-element only needs to contain ORFs encoding LOV2 and Zdk1 and the required regulatory elements, while the flanking homology arms direct it to the correct genomic locus. The rest of the protein of interest remains encoded by the endogenous gene. Here, we also include a fluorescent protein tag to monitor π-EB1 photodissociation and a leucine zipper coiled-coil to retain dimerization of the π-EB1 N-terminal half during blue light exposure. While we demonstrate the utility of this approach for the MAPRE1 gene in human iPSCs, we believe that a similar strategy could be adapted to many other multidomain proteins supplementing the optogenetic toolbox to investigate localized protein functions in real time (Wittmann et al., 2020), but it should be noted that photodissociation kinetics may be different with monomeric proteins. The LOV2/Zdk1 module interacts in the dark, which is opposite to all other optogenetic dimerization modules and therefore enables an unperturbed dark state and acute functional knockout of a given protein activity through blue light exposure. Thus, replacing an endogenous protein with a photosensitive variant in a single genome editing step as presented here has important advantages by allowing modification of proteins for which genetic knockouts might be lethal and further enabling optogenetics in complex cell systems that are not amenable to transient genetic manipulations.

This approach allowed us to directly demonstrate that EB1 in developing human cortical neurons is required for sustained growth cone MT growth and growth cone advance. Although we do not completely dissect the molecular mechanism underlying the π-EB1 photodissociation-mediated neurite retraction response, unchanged f-actin polymerization dynamics during blue light indicate that retraction likely results from a loss of growth cone adhesion or an increase in neurite actomyosin contractility consistent with MT-shortening induced RhoA activation (Joo and Olson, 2021). An increase in neurite contractility that does not remain localized to the region of blue light exposure may also explain our difficulty in using π-EB1 photodissociation to control growth cone guidance more precisely. Thus, although other optogenetic tools that provide attractive stimuli could be a promising avenue to control growth cone guidance (Harris et al., 2020), inhibitory stimuli that induce MT depolymerization may be too difficult to control to be practically useful in synthetic biology approaches building defined neuron networks. Nevertheless, it will be interesting to see how π-EB1 in i^3^N or other types of neurons can be used to analyze how MTs and +TIPs contribute dynamically to other aspects of neuron morphogenesis such as branching, dendritic spine dynamics or synaptic plasticity (Dent et al., 2004; Jaworski et al., 2009)

## Materials and Methods

### Molecular cloning and genome editing

#### Construction of knock-in π-EB1 cassettes

Primer sequences for all cloning, gRNAs and genomic PCR are given in supplemental Table S1, and relevant plasmids will be made available in Addgene.

1. pEB1N-LZ-LOV2-IRES-mCherry-Zdk1-EB1C was cloned as follows: EB1N-LZ-LOV2 (van Haren et al., 2018) (Addgene plasmid #107614) and EMCV IRES (from MSCV-PIG; Addgene plasmid #18751) were amplified by PCR, connected by overlap extension PCR, and ligated into the NheI site of pmCherry-Zdk1-EB1C (Addgene plasmid #107695).

2. pUC19-π-EB1_IRES-mCherry (LZ-LOV2-IRES-mCherry-Zdk1) containing the homology-directed repair (HDR) template for targeting the IRES containing version of the π-element into exon 5 of MAPRE1/EB1 was cloned as follows: >1.5 kb 5’and 3’ homology arms were amplified from genomic DNA isolated from one 6-cm dish of immortalized human retinal pigment epithelial cells (hTERT-RPE1 cells, ATCC) using a Purelink Genomic DNA Mini Kit (ThermoFisher). The π-element was PCR amplified from plasmid #1. Homology arms and π-element sequences were cloned into the EcoRI site of pUC19 by Gibson assembly in a single step. Several silent mutations were introduced to ensure that the HDR template could not be targeted by the gRNAs.

3. pUC19-sv40-blastR-EF1α-mCherry-Zdk1-EB1C was generated by Gibson assembly of PCR amplified SV40promoter-BlasticidinR, SV40polyA, EF1alpha promoter (from a PiggyBac vector) and mCherry-Zdk1-EB1C-polyA (from pmCherry-Zdk1-EB1C) into HinDIII and EcoRI sites of pUC19.

4. pEB1N-LZ-LOV2-polyA-EF1alpha-mCherry-Zdk1-EB1C was generated by Gibson assembly of PCR amplified polyA-EF1alpha-mCherry-Zdk1-EB1C from plasmid #3 into the BamHI site of pEB1N-LZ-LOV2.

5. pUC19-π-EB1_EF1alpha-mCherry (LZ-LOV2-polyA-EF1α-mCherry-Zdk1) containing the HDR template for targeting the π-element to exon 5 of MAPRE1/EB1 was cloned by Gibson assembly of PCR amplified homology arms and π-element into the EcoRI site of pUC19 using the same primers as for the IRES plasmid (#2).

6. For generating π-EB1 knock-in hiPSCs an EGFP-tagged variant of the π-element was used. pUC19-π-EB1_EF1α-EGFP was generated by replacing mCherry in plasmid #5 with EGFP by Gibson assembly of PCR-amplified EGFP into the NcoI sites.

#### Guide RNAs

7. pSpCas9(BB)-2A-GFP was a gift from Feng Zhang (Addgene plasmid #48138). pSpCas9(BB)-2A-mCherry was constructed by replacing GFP with mCherry. T2A-mCherry was PCR amplified and inserted into the EcoRI sites by Gibson assembly.

8. Three gRNA sequences that target <25 bp from the integration site in exon 5 of MAPRE1/EB1 were cloned into the BbsI sites of pSpCas9(BB)-2A-GFP (Wittmann and van Haren, 2018) to knock in the π-element into the EB1 gene.

9. Similarly, three gRNA sequences targeting exon 1 of MAPRE3 were cloned into the BbsI sites of pSpCas9(BB)-2A-GFP and pSpCas9(BB)-2A-mCherry to generate EB3 knockout lines.

#### Other DNA constructs

10. EB1N-mScarlet-I-LZ-LOV2 and EB1N-mApple-LZ-LOV2 were cloned by inserting PCR amplified mScarlet-I/mApple and LZ-LOV2 into the XhoI and BamHI sites of EB1-GFP-LOV2 by Gibson assembly. While the original EB1N-mCherry-LZ-LOV2 fusion construct did not perform well in iPSC derived neurons (and showed striking aggregation), the mScarlet and mApple fusions showed the expected localization pattern at MT plus-ends. The two resulting PCR products were inserted into the XhoI/BamHI sites of plasmid EB1N-GFP-LOV2. mScarlet-I was PCR amplified using an mRuby2 forward oligo, which resulted in a slightly altered mScarlet-I N-terminus (MVSKGEEL instead of MVSKGEAV).

#### i^3^N cell culture and differentiation

i^3^N hiPSCs were cultured essentially as described (Wang et al., 2017). In brief, stem cells were cultured on Matrigel-coated dishes with mTeSR medium changed every 2 days for 3-5 days, until 70-80% confluent, then gently detached with Accutase and reseeded as 2-5×10^4^ cells/cm^2^ in mTeSR medium containing 10uM Y-27632 (#72304, STEMCELL Technologies) ROCK inhibitor (removed after 48h and exchanged for fresh medium without ROCK inhibitor). The stem cells were transfected with Lipofectamine Stem (STEM00015, Invitrogen) 24h after seeding according to manufacturer’s protocol.

For pre-differentiation, 2-3×10^6^ i^3^N stem cells were seeded on a Matrigel-coated 3.5 cm dish (high crowding helps the early steps of the differentiation process) in KO DMEM supplemented with 1x N2 supplement, 2 µg/ml doxycycline, 10 ng/ml recombinant neurotrophin-3 (NT-3), 10 ng/ml recombinant brain-derived neurotrophic factor (BDNF), 0.3 µg/ml murine laminin and 10uM Y-28632 ROCK inhibitor (only for the first day, then withdrawn). The medium was changed every 24h for 3 days of culture.

For imaging-directed differentiation, 20-mm glass-bottom Mattek dishes were surface-activated for 1-2 minutes in a Harrick Expanded Plasma Cleaner (Dema et al., 2022b) set on “High” and coated with 50 µg/ml poly-D-lysine (PDL) in PBS for 20 minutes at 37°C, washed extensively in PBS, then coated with 1.5 µg/ml murine laminin in PBS for 20 minutes at 37°C. After a further PBS wash, 2×10^4^ to 10^5^ pre-differentiated i^3^N cells from the Accutase-dissociated cell pellet were seeded in Maturation Medium (MM), consisting of 50% Neurobasal and 50% DMEM-F12 supplemented with 1x N2 supplement, 1x B27 supplement, 2 µg/ml doxycycline, 10 ng/ml NT-3, 10 ng/ml BDNF and 0.3 µg/ml murine laminin. Laminin micropatterns on glass-bottom dishes were generated as described (Dema et al., 2022b). For immunoblots or qPCR, the pre-differentiated cells were seeded at 10^5^ cm^-1^ on PDL and laminin-coated tissue culture plastic dishes. Cell lysate preparation and immunoblotting for EB1 and EB3 were performed as described (van Haren et al., 2018). Mouse monoclonal anti-KIF2C antibody (2488C3a) was from Santa Cruz Biotechnology (SC-81305).

#### Genome editing

In general, cells were co-transfected with two or three gRNA Cas9 plasmids targeting MAPRE1/EB1 exon 5 and a pUC19 HDR template described above. H1299 cells were transfected using Lipofectamine 2000 in a 10-cm dish at ∼75% confluency and 2-3 days later mCherry-positive cells (indicating π-element integration) were FACS sorted into 96-well plates for expansion and further analysis.

For generating i^3^N π-element lines, both the π-element and gRNA plasmids carried EGFP markers. Thus, cells were FACS sorted 6 days after transfection when we expected that most cells had lost transient expression from the gRNA plasmids. EGFP-positive i^3^N cells were then plated at low density (∼3000 cells) in 6-cm Matrigel coated dishes and cultured in mTeSR with ROCK inhibitor as described above. After 7-10 days, clonal colonies typically had grown enough to be seen by naked eye. Up to 96 colonies from each dish were manually picked and expanded. Initial screening for positive clones in both H1299 and i^3^N cells was by spinning disk confocal microscopy because clones in which fluorescent signal was present on MT plus ends in the absence of blue light must have correctly integrated the π-element such that both halves are expressed as the N-terminal half (π-EB1N) expressed from the endogenous promoter is needed to target EGFP-tagged π-EB1C to MT plus ends. Together with wildtype controls, such clones were further analyzed by genomic PCR with primer sets that bind inside and outside the π-element sequence (supplemental Table S1). Genomic DNA was isolated using a Purelink Genomic DNA Mini Kit (ThermoFisher). Expected PCR products for EGFP EF1α π-element: primer pair 11a/11b, WT: 189 bp, π-element integration: 3058 bp; 11c/11b, WT: 1628 bp; π-element integration: 4506 bp; 11e/11f, WT: no product, π-element integration: 2152 bp.

For the MAPRE3/EB3 knockout in π-element i^3^N cells, gRNAs were cloned into an mCherry version of the Cas9 plasmid, and transfected cells FACS-sorted for mCherry expression, plated at low density and clones expanded as described above. Initial positive clones identified by band shift in genomic PCR (see supplemental Table S1 for primer sequences) were further analyzed by Sanger sequencing of the PCR amplicon and analyzed by ICE (Synthego Performance Analysis, ICE Analysis. 2019. v3.0. Synthego).

#### Live microscopy and π-EB1 photoinactivation

i^3^N neuron imaging experiments were performed 1-3 days after replating on laminin-coated glass-bottom dishes. At later times i^3^N neurons form intricate networks and it becomes increasingly difficult to locate individual growth cones. SPY555 tubulin and SPY650-FastAct (Spyrochrome) were added to i^3^N neurons at a 1:2000 and 1:3000 dilution, respectively, and 4-610CP-CTX was added at 5 nM for at least 30 min before imaging, and cells were discarded after a maximum of 3 hours.

Live microscopy was performed either with a Yokogawa CSU-X1 spinning disk confocal essentially as described (Stehbens et al., 2012) or, for most i^3^N neuron microscopy, with a CFI Apochromat TIRF 60X NA 1.49 objective (Nikon) on a Yokogawa CSU-W1/SoRa spinning disk confocal system, and images acquired with an ORCA Fusion BT sCMOS camera (Hamamatsu).

For high-resolution imaging of dim signal, SoRa mode was combined with 2×2 camera binning resulting in an image pixel size of 54 nm. This system was equipped with a Polygon 1000 pattern illuminator (Mightex) through an auxiliary filter turret and LAPP illuminator (Nikon). Integrated control of imaging and photoinactivation was through NIS Elements v5.3 software (Nikon), in combination with an external pulse generator to trigger the 470 nm photoinactivation LED (Spectra X light engine, Lumencor) for 10 ms pulses at 2 Hz at 5-10% LED power. π-EB1 photoinactivation was confirmed by dissociation of the C-terminal half from growing MT ends. The Polygon was calibrated with a mirror slide before every imaging session, and local light exposure further verified by imaging the back reflection from the specimen coverslip.

#### Image analysis and statistics

MT growth rate analysis in i^3^N hiPSCs was as described. MT ends in SPY555-tubulin labeled i^3^N neurons as well as neurite length changes were tracked manually using the ‘Segmented Line’ tool of FiJi. Kymographs to measure retrograde F-actin flow were generated as described (Stehbens et al., 2014). Details of statistical analysis including the type of test, p-values and numbers of biological replicates are provided within the relevant figures and figure legends. All statistical analysis was done in MATLAB (Mathworks, Inc.)., and graphs were produced in MATLAB and in Excel (Microsoft). In all figures, box plots show median, first and third quartile, with whiskers extending to observations within 1.5 times the interquartile range.

## Acknowledgements

We thank Li Gan for the i^3^N hiPSCs, and Gražvydas Lukinaviĉius and Jonas Buceviĉius for the generous gift of fluorogenic cabazitaxel MT probes. This work was supported by National Institutes of Health grants R21 CA224194, R01 NS107480, S10 RR026758 and S10 OD028611 to T.W.

