## Supplemental Material for "Growth cone advance requires EB1 as revealed by genomic replacement with a light-sensitive variant"

**Supplementary figures**

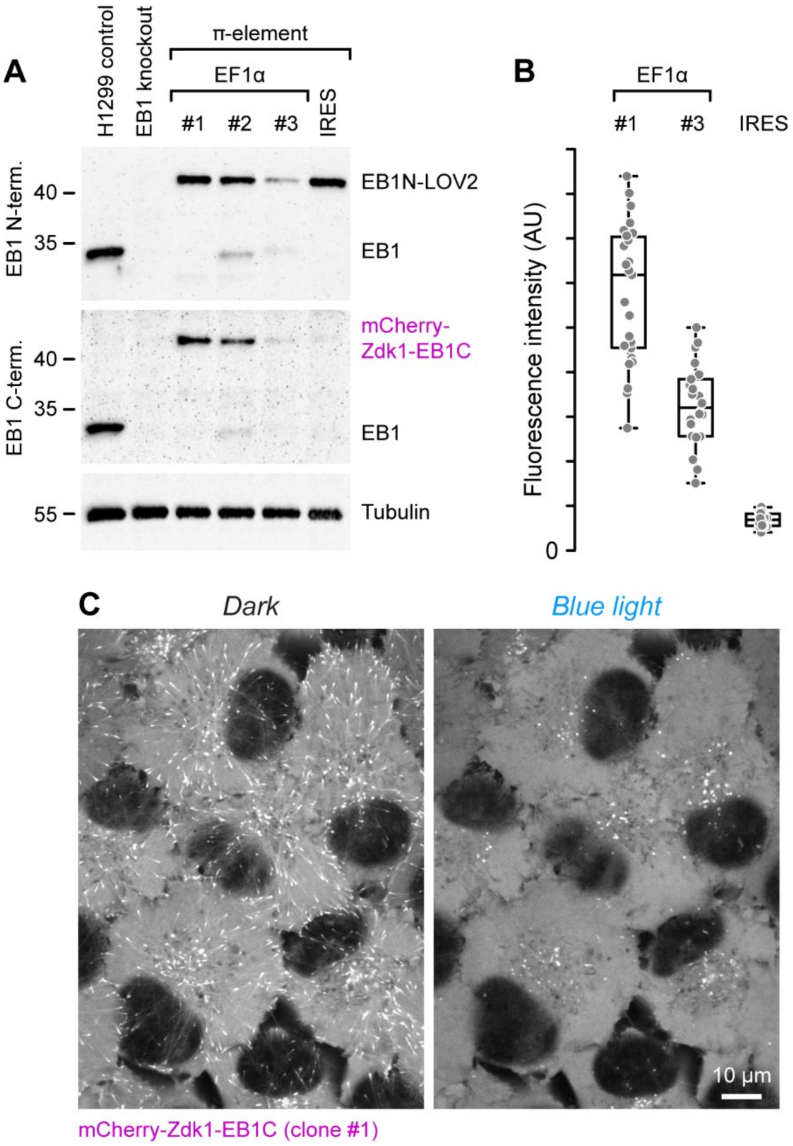


**Figure S1. Evaluation of π-element designs in H1299 lung cancer cells. (A)** Immunoblot analysis of EB1 expression in cells in which different π-element designs were inserted into the MAPRE1/EB1 gene. EF1α clone #1 likely represents a homozygous insertion with near endogenous expression levels of both π-EB1 parts indicating that the MAPRE1/EB1 and EF1α promoters are well matched, while clones #2 and #3 have remaining wild-type EB1 expression. Note that the IRES π-element (EMCV-IRES with A7 bifurcation loop) has nearly no expression of the C-terminal π-EB1 half, while expression of the N-terminal half is nearly identical to EF1α clone #1. This is consistent with previous reports showing that IRES-dependent expression of the second gene is often much lower than cap-dependent first gene expression but might be improved with different IRES variants (Bochkov and Palmenberg, 2006). **(B)** Fluorescence intensity of mCherry-Zdk1-EB1C in the indicated clones again showing very low expression in cells with the IRES π-element. Each data point represents the mean fluorescence intensity of one cell. **(C)** Images of EF1α clone #1 showing dissociation of mCherry-Zdk1-EB1C from growing MT ends during blue light exposure.

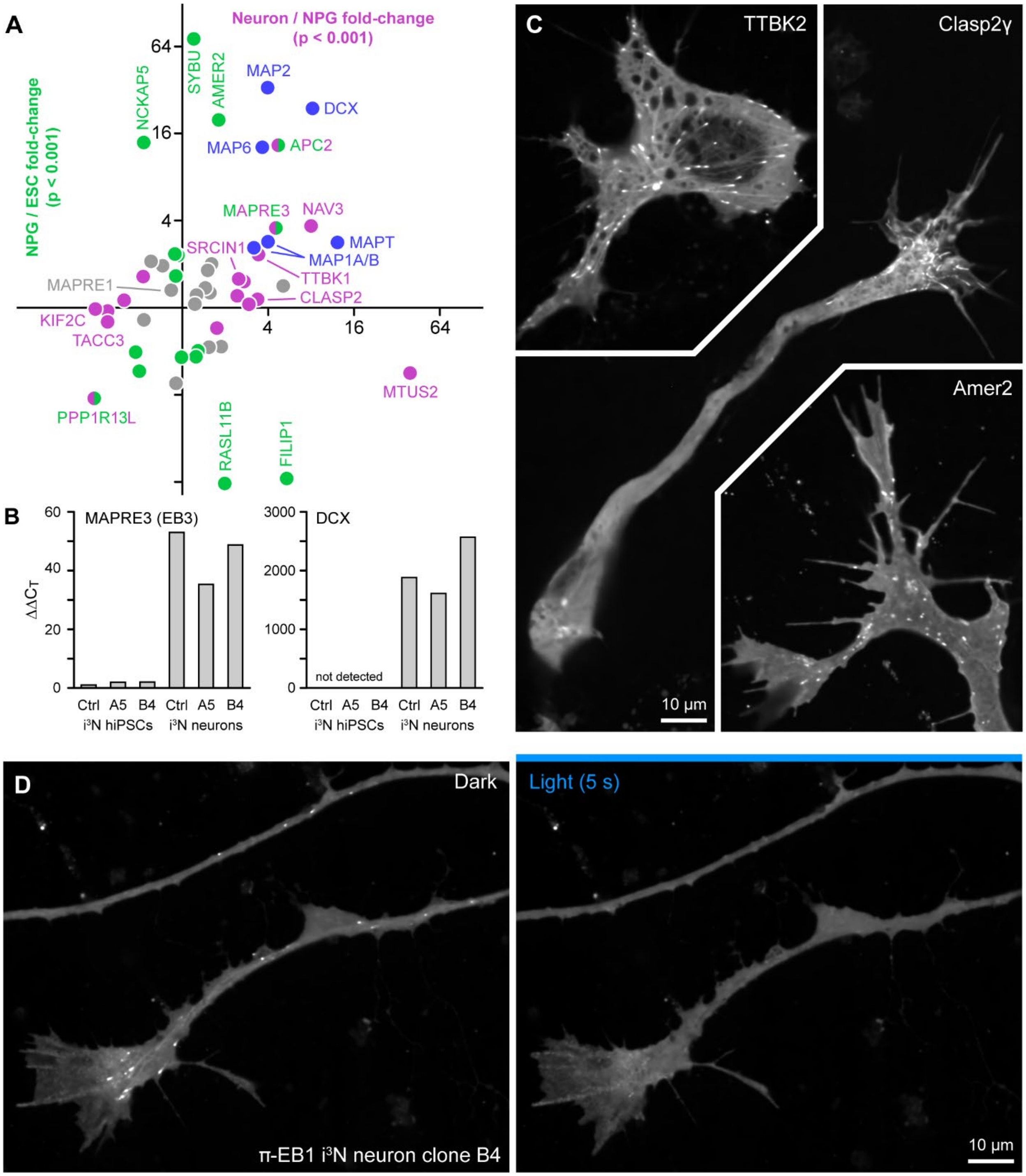


**Figure S2. Gene expression changes associated with neuron differentiation. (A)** Changes of MAP and +TIP translation during human neuron differentiation based on a published ribosome profiling data set (Blair et al., 2017). The vertical axis indicates changes between embryonic stem cells (ESCs) and neuron progenitor cells (NPGs) and the horizontal axis changes between

NPGs and neurons. Changes of colored symbols were statistically significant to a $p < 0.001$. Selected proteins are labelled. **(B)** RT-qPCR analysis of EB3 and DCX expression in control and π-EB1 $i^3N$ lines illustrating robust upregulation of both proteins during Ngn2-induced neuronal differentiation. **(C)** Examples of growing MT plus end localization of transiently transfected EGFP-tagged +TIPs in $i^3N$ neurons in early stages of differentiation. Note that Clasp2γ preferentially binds growth cone MTs likely due to a GSK3β kinase activity gradient (Kumar et al., 2009). Amer2 also displays prominent membrane localization. **(D)** π-EB1 $i^3N$ neurite and growth cone showing dissociation of the EGFP tagged π-EB1 C-terminal half during blue light exposure.

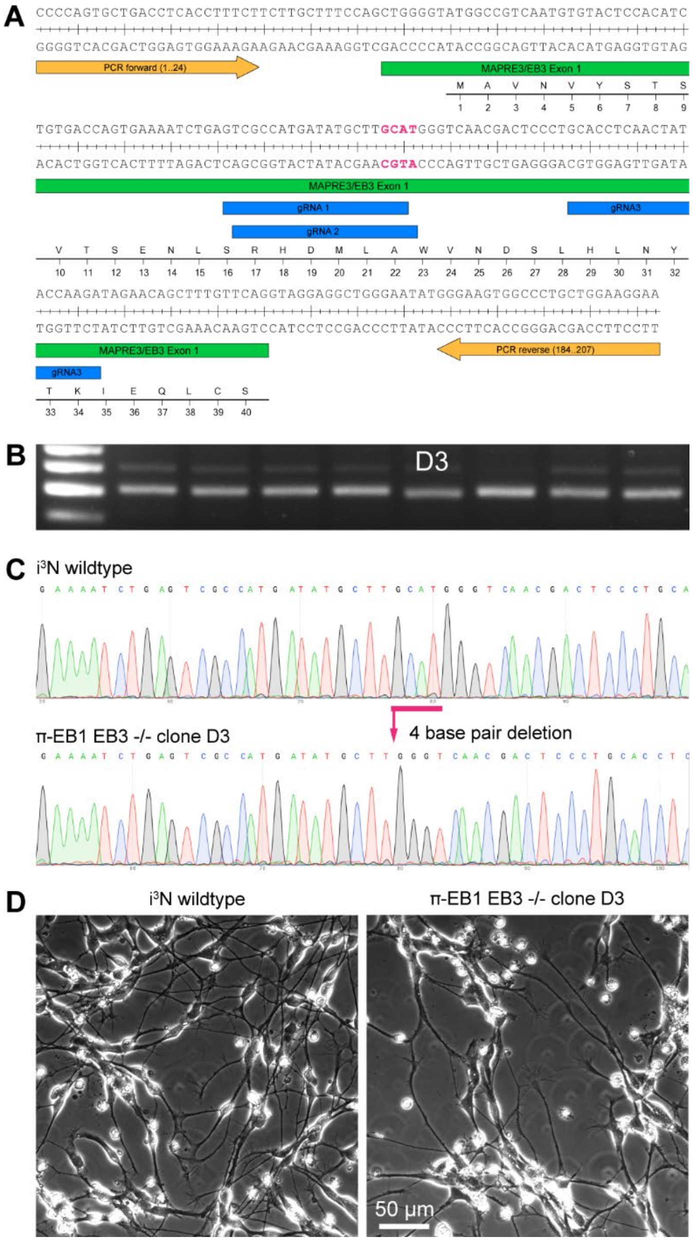


**Figure S3. Validation of EB3 knockout in π-EB1 i$^3$N cells. (A)** MAPRE3/EB3 knockout strategy by CRISPR/Cas9 genome editing. **(B)** Genomic PCR across MAPRE1/EB1 exon 1 with the primers indicated in (A) showing a small downshift in the clone labeled D3. **(C)** Sequencing of the genomic PCR products from EB3 wild-type cells and the knockout clone indicates a four base pair deletion, also highlighted in red in (A) that introduces a frame shift and stop codon in exon 1. **(D)** Images of wildtype and π-EB1 EB3-/- i$^3$N neurons showing normal differentiation two days after plating on laminin-coated dishes.

**Supplementary video legends**

**Video 1.** Time-lapse of genome edited π-EB1 $i^3N$ hiPSCs transiently expressing an EB1N-mScarlet to visualize MT growth dynamics before and during blue light exposure. Related to Fig. 1G.

**Video 2.** Long-term phase contrast time-lapse of π-EB1 EB3-/- $i^3N$ neurons in the absence of blue light exposure showing normal neurite and growth cone dynamics. The cells are growing on a grid of laminin paths. Elapsed time is shown in hours:minutes:seconds.

**Video 3.** Time-lapse of EGFP-tagged π-EB1 C-terminal half in π-EB1 EB3-/- $i^3N$ neurons demonstrating repeatability of π-EB1 photodissociation in differentiating neurons. 3-second photoinactivation sequences are interspersed by 5 min dark recovery periods. Elapsed time is shown in minutes:seconds. Related to Fig. 2A.

**Video 4.** Time-lapse of MT dynamics in SPY555-tubulin labeled control and π-EB1 EB3-/- $i^3N$ neuron growth cones before and during blue light exposure. Related to Fig. 2B.

**Video 5.** Time-lapse of a growth cone of a π-EB1 EB3-/- $i^3N$ neuron labeled with SPY650-FastAct before and during blue light exposure indicating that f-actin retrograde flow is insensitive to π-EB1 photodissociation at this time scale. Related to Fig. 3A.

**Video 6.** Time-lapse of control and π-EB1 EB3-/- $i^3N$ neurite dynamics in which MTs were labeled with 4-610CP-CTX before and during blue light exposure. Related to Fig. 4A.

**Video 7.** Long-term phase contrast time-lapse of a π-EB1 neurite advancing along laminin path toward a blue light exposed region. The growth cone fails to cross the blue light barrier multiple times. Related to Fig. 4C. Elapsed time is shown in hours:minutes:seconds.
